# Ground-truth evaluation of sparse autoencoder features on spatial, single-cell and language-model embeddings

**DOI:** 10.64898/2026.10.27.762296

**Authors:** Amirhossein Shahsavand, Shayan Forghani, Saba Akbarzadeh, Fatemeh Mahdavi Sabet

## Abstract

Sparse autoencoders (SAEs) are increasingly used to extract interpretable features from the embeddings of biological foundation models. Because SAEs trained with different random seeds learn different features, reproducibility across seeds has become a common test of whether these features can be trusted. We asked whether reproducible features are also meaningful, using embeddings whose concepts are known: five spatial transcriptomics embeddings of two colorectal cancer sections and Geneformer embeddings of single-cell RNA profiles, with cell lineages as concepts; synthetic data with planted cell states; and the residual stream of a small language model, with token classes as concepts. Varimax rotations of the same embeddings were almost perfectly reproducible, yet rotation features (PCA, varimax and ICA) covered at most three of seven lineages and three of eight token classes. Single SAE fits showed the opposite profile: they covered most concepts with purer gene programs but reproduced only 27–46% of their features. A consensus over five SAE fits combined the two strengths. It was reproducible (78–98% across sources) and as pure as single fits: 11–40% of its features carried marker genes of another lineage, against 44–70% for varimax. Against the strongest baseline, varimax sparsified to the SAE’s sparsity, the consensus covered as many concepts on PCA and Geneformer embeddings and more on learned spatial and text embeddings. The consensus also had costs. It covered fewer lineages than single fits and, like all SAEs, missed weak planted states that rotations recovered; neither a longer single fit nor a second SAE trained on the consensus residual recovered what it discarded. Reproducibility alone therefore ranks features poorly: SAE features should be reported with the number of known concepts they cover and the purity of their gene programs, against a rotation of the same embedding sparsified to the SAE’s sparsity.

## Introduction

Foundation models for single-cell transcriptomics^1,2^ and representation methods for spatial transcriptomics^3^ compress each cell into a dense embedding of a few dozen to a few thousand dimensions. These embeddings support clustering, annotation and transfer across datasets, but their individual dimensions rarely correspond to recognizable biology. Sparse autoencoders (SAEs) offer a way to read them. An SAE re-expresses each embedding as a sparse, non-negative combination of a larger set of learned directions, and each direction is then inspected as a candidate concept^4,5^. SAEs developed for language models^6,7^ are now applied to protein language models^8^ and to single-cell foundation models^9^.

Two problems limit how far SAE features can be trusted. The first is that SAEs trained on the same data with different random seeds learn substantially different features^10^. A feature reported from one fit may simply not exist in another. The second is that interpretability is usually judged without ground truth: simple baselines often match SAEs on downstream tasks^11,12^, and SAEs with random components can score as well as trained ones on common interpretability metrics^13^. Recent work addresses the first problem by making features agree across seeds, through aligned encoder and decoder directions^14^, archetypal constraints^15^, or pooling of features across seeds^16^. Each of these methods reports reproducibility as a main measure of success. Whether reproducibility indicates that features are meaningful has not been tested directly. A follow-up study traced the stability of archetypal SAEs to shared initialization rather than to the learned dictionary^17^, which illustrates the risk.

Here we tested reproducibility against two properties that matter for biological interpretation. The first is coverage: how many known concepts the features represent. The second is purity: whether each feature represents one concept rather than a mixture. We used embeddings where concepts are known. Cell lineages are annotated independently of any embedding, cell states can be planted in synthetic data, and token classes in text can be defined by rules. Our main comparison is with rotations of the same embedding. A rotation such as varimax^18^ is fully determined by the data, so it is reproducible by construction, but it has no reason to align with concepts. Because dense rotation components are active in most cells, we also sparsified varimax to the SAE’s number of active features per cell, which gives rotations the same opportunity to become concept-specific.

We asked three questions. Do SAE features cover concepts better, and more purely, than rotations? Does a consensus over seeds, analogous to consensus non-negative matrix factorization (cNMF)^19^, make SAE features reproducible, and what does it discard? Can a longer single fit, or a second SAE trained on what the consensus misses, recover the discarded information? We answered these questions on five spatial embeddings, planted synthetic data, Geneformer embeddings and a language model, with three seeds each.

## Results

### A test bench with known concepts

To ask whether reproducible SAE features are also meaningful, we needed embeddings whose meaningful content is known in advance. We assembled four such sources (Methods). The first was two colorectal cancer sections profiled by CosMx spatial molecular imaging^20^ with the whole-transcriptome panel^21^: a section from a study of colorectal malignant transformation^22^ (131,000 cells) and a public colon dataset (494,000 cells). We analyzed 50,000 cells per section on a fixed set of 2,000 genes and embedded each section with three methods: principal components of log-normalized expression (Expr50), CellCharter^3^ and, for the second section, CadaST^23^. This gave five embeddings of 50–256 dimensions, in which the known concepts were seven cell lineages assigned by reference-based cell typing^24,25^, independently of any embedding. The second source was embeddings from the Geneformer V2 model^2^ (1,152 dimensions) of 12,881 cells from a colorectal cancer atlas^25^, with the same seven lineages. Lineages are strong signals, so we added a synthetic dataset in which weaker biology is known exactly: six lineages, each with three cell states of decreasing strength, and four programs shared across lineages (28 programs in total), simulated as counts and embedded like Expr50. The fourth source, outside biology, was the residual stream of the Pythia-70m language model^26^ (512 dimensions) on one million WikiText-103 tokens^27^, with eight rule-defined token classes, such as numbers, punctuation and function words, as concepts.

On each embedding we trained a BatchTopK SAE^28^, which describes each cell by a small number of active features, both as single fits and as a consensus of five fits with different seeds. As baselines we used rotations, which re-express the same embedding along new axes without learning a dictionary: PCA, varimax, independent component analysis (ICA)^29^ and a sparse varimax that keeps, for each cell, only as many active features as the SAE uses. We scored every set of features in three ways. Reproducibility (R1) is the share of features that reappear, with activation correlation r ≥ 0.7, when the method is refitted on the other half of the cells. Coverage is the number of known concepts represented by at least one concept-specific feature, that is, a feature with at least 70% of its activity in one concept; on planted data it is the number of programs matched at r ≥ 0.7. Purity is judged by the share of chimeric features, whose top genes include several markers of other lineages. All values are means over three seeds.

### SAE features cover more concepts than rotations, with purer gene programs

We first asked which methods represent the known concepts. Dense rotations rarely did (Fig. 1, Table 1). On the spatial embeddings, varimax, PCA and ICA covered at most two of seven lineages; on Geneformer they covered at most three, and on the language model at most three of eight token classes. SAE features did much better. The consensus covered 2–5 lineages on the spatial embeddings and every concept on Geneformer (7 of 7) and on the language model (8 of 8), and single fits covered up to 6.7 lineages.

**Table 1.** Consensus SAE against each method. For the spatial embeddings and synthetic settings, wins/ties/losses of the consensus, where a difference counts only if it exceeds the sum of the seed standard deviations. For Geneformer and the language model, values are shown directly (consensus vs comparison). Coverage is lineages (of 7), planted programs (of 28) or token classes (of 8). Higher is better except for chimeric features.

| Source | Comparison | Coverage | Chimeric features | Hallmark | kNN | R1 |
| --- | --- | --- | --- | --- | --- | --- |
| Spatial (5) | sparse varimax | 3/1/1 | 5/0/0 | 5/0/0 | 3/1/1 | 0/1/4 |
| Spatial (5) | varimax | 5/0/0 | 5/0/0 | 4/0/1 | 2/1/2 | 0/1/4 |
| Spatial (5) | ICA | 5/0/0 | 5/0/0 | 4/0/1 | 5/0/0 | 3/2/0 |
| Spatial (5) | single SAE | 0/2/3 | 2/1/2 | 1/3/1 | 3/2/0 | 5/0/0 |
| Synthetic (3) | sparse varimax | 2/0/1 | – | – | – | 0/0/3 |
| Synthetic (3) | varimax | 1/2/0 | – | – | – | 0/0/3 |
| Synthetic (3) | single SAE | 1/2/0 | – | – | – | 3/0/0 |
| Geneformer | sparse varimax | 7 vs 7 | 0.09 vs 0.27 | 0.72 vs 0.60 | 0.88 vs 0.74 | 0.85 vs 0.67 |
| Geneformer | varimax | 7 vs 0 | 0.09 vs 0.47 | 0.72 vs 0.61 | 0.88 vs 0.86 | 0.85 vs 1.00 |
| Language model | sparse varimax | 8 vs 6 | – | – | 0.89 vs 0.73 | 0.78 vs 1.00 |
| Language model | varimax | 8 vs 1 | – | – | 0.89 vs 0.91 | 0.78 vs 1.00 |

**Figure 1.**
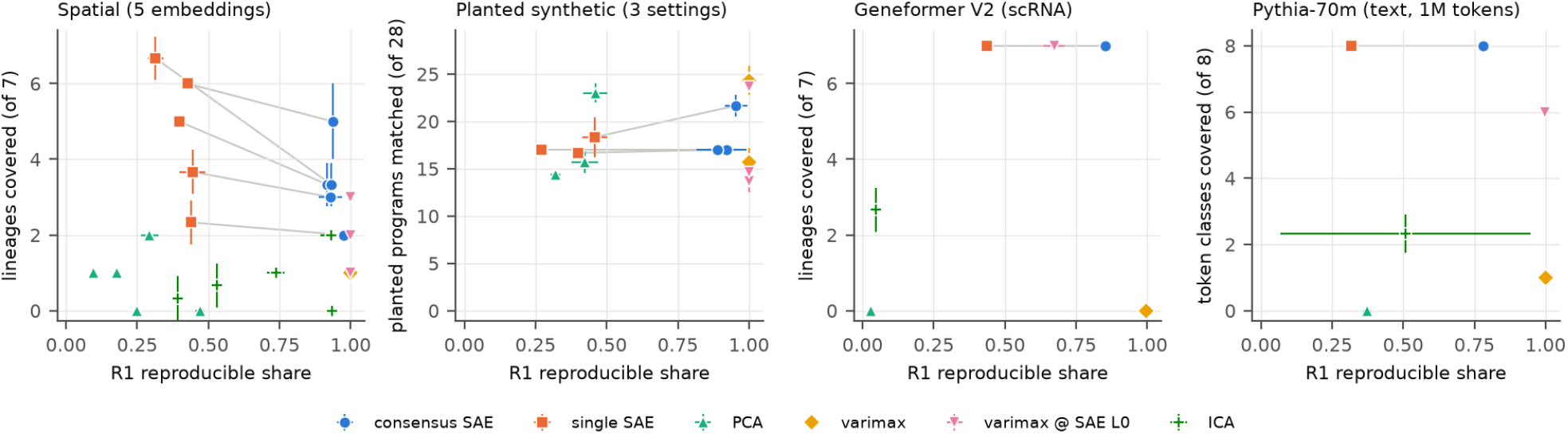
Reproducibility and concept coverage of SAE and rotation features. Reproducibility (R1, share of features recurring between models fitted on random halves) against the number of distinct concepts covered, for each embedding and method. Panels, left to right: five spatial embeddings (lineages of 7); planted synthetic data (programs of 28 matched at r ≥ 0.7); Geneformer embeddings (lineages of 7); Pythia-70m residual stream, one million tokens (token classes of 8). Gray lines join the single and consensus SAE of the same embedding. Points are means and bars standard deviations over three seeds. On the real embeddings, varimax reaches R1 ≈ 1 with almost no coverage and PCA and ICA are less reproducible; single SAE fits cover most concepts at low R1; the consensus SAE raises R1 at some loss of coverage.

Part of this advantage comes from sparsity alone. A dense rotation axis is active in most cells and so is rarely specific to one lineage, whereas a sparse one can be. Sparse varimax indeed closed the gap on the Expr50 embeddings (3 lineages, against 2–3 for the consensus) and on Geneformer (7 against 7), but not on the learned CellCharter and CadaST embeddings (1–2 against 3.3–5) or on text (6 against 8 classes). Across the five spatial embeddings, the consensus covered more lineages than sparse varimax on three, equal on one and fewer on one.

Purity separated the methods more clearly than coverage. For each feature we listed the genes whose expression rises with its activity; a feature whose list includes several markers of other lineages mixes cell types. Such chimeric features made up 11–40% of the consensus, against 44–70% for varimax and 46–69% for sparse varimax, and the consensus was purer than every rotation on all five spatial embeddings. On Geneformer, 9% of consensus features were chimeric, against 47% for varimax. Consensus features were also more often enriched for an MSigDB Hallmark gene set^30^ than sparse-varimax features (5 of 5 embeddings).

Planted data, where every program is known, put these gains in proportion. SAEs did not recover more planted programs than rotations: the consensus matched 17–22 of 28 programs, single fits 17–18, varimax 16–24 and PCA 14–23. The methods differed instead in how they used their features. Almost all consensus features (91–94%) matched a planted program, but many were copies of the same strong lineage identity, 2.9–5.3 per lineage against 1.3–1.6 for rotations. Most rotation features (about 80 of 100) matched nothing, because half of each split component is a negative direction and many components carry only noise. Because of these copies, counts of concept-specific features, which reached 217 on the spatial embeddings and 418 on text, overstate how many concepts an SAE finds, and we report coverage instead.

### Reproducibility ranks the methods in reverse

Judged by reproducibility, the ranking reversed (Fig. 1). Single SAE fits reproduced only 27–46% of their features when refitted on the other half of the data, whereas varimax reproduced 99–100% on every source. Yet varimax covered at most one concept on any biological or text embedding.

### A consensus over seeds keeps the reproducible, pure features

To obtain reproducible SAE features, we combined five fits in the spirit of consensus non-negative matrix factorization (cNMF)^19^: features that match across fits are grouped, with at most one feature per fit, and a group is kept if it appears in at least three of the five fits (Methods). The consensus raised reproducibility to 92–98% on the spatial embeddings (above single fits on all five), 89–95% on planted data, 85% on Geneformer and 78% on the language model (Fig. 2, left). In absolute numbers it held only modestly more reproducible features than a single fit (5–27% more; higher beyond the seed variation on 2 of 5 spatial embeddings, Geneformer and the language model, and tied on the other three; Fig. 2, middle). Its main gain was therefore to identify which features recur, so that each retained feature can be interpreted with confidence. Its features were as pure as those of single fits (chimeric share lower on 2, tied on 1 and higher on 2 of 5 embeddings). A consensus anchored on the first fit gave similar results on the real embeddings but was less reproducible on planted data (0.85 against 0.95 at depth 500; Supplementary Table S5).

**Figure 2.**
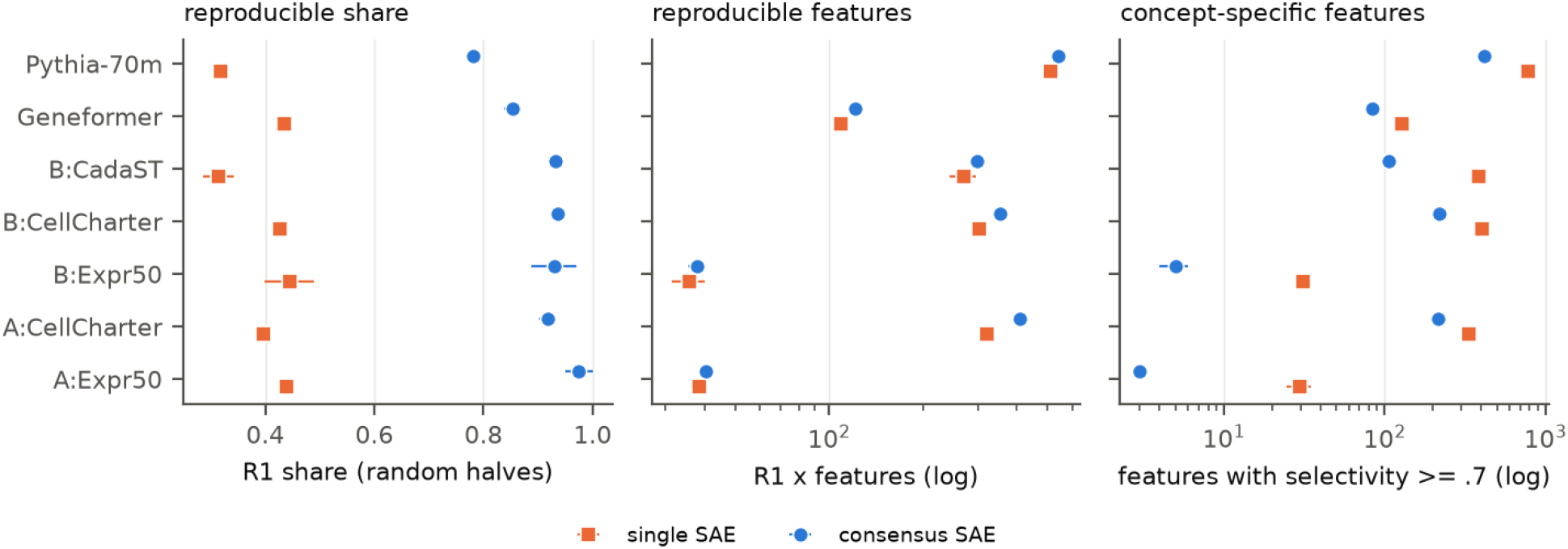
The consensus as a reliability filter. Single and consensus SAEs on each spatial, Geneformer and language-model embedding. Left, reproducible share (R1). Middle, number of reproducible features (R1 × features, log scale). Right, number of concept-specific features (log scale); these counts include many duplicates of the same concept. The consensus raises the reproducible share, with only a modest increase in the number of reproducible features. Means and standard deviations over three seeds.

### The dictionary must be bounded by the number of cells

A reliable consensus also required a dictionary that is not too large for the data. If an SAE has many more latent features than the data can support, each fit can divide the data differently. We therefore used four latents per embedding dimension, but no more than about one latent per 50 cells or tokens (Methods). This cap applied only to Geneformer, where 12,881 cells allow about 257 latents instead of 4,608; with 256 latents, the consensus reached 85% reproducibility. Larger dictionaries lowered reproducibility steadily (Matryoshka consensus on Geneformer: 0.80, 0.60 and 0.25 with 256, 1,152 and 4,608 latents). On the language model, more data helped: the consensus reached 0.62 with 50,000 tokens (Matryoshka) and 0.78 with one million (plain BatchTopK), but stayed below the biological embeddings. This difference follows the geometry of the embeddings, to which we return in the Discussion. The language-model embedding spreads its variance over about 70 effective dimensions, against 8.5 for Geneformer and 10–20 for the spatial embeddings, and its token classes explain about half as much variance as lineages do (Supplementary Table S4).

### What the consensus discards

The consensus came at a cost (Fig. 1; Fig. 2, right). Compared with single fits, it covered fewer lineages on 3 of 5 spatial embeddings (median 1.0 fewer) and kept 7–64% of their concept-specific features. Most of the discarded features were copies of retained concepts, but some lineages were lost entirely. The consensus also needed five fits, four to five times the run time of a single fit and 30 to 300 times that of a rotation. Its features separated fine cell types as well as varimax on the spatial and Geneformer embeddings (kNN accuracy) and slightly less well on the language model (0.89 against 0.91).

The largest gap concerned weak cell states, and it was shared by all SAEs rather than caused by the consensus (Fig. 3). At sequencing depth 500, single fits recovered 0.89, 0.33 and 0.17 of the strong, mid and weak states, the consensus 1.00, 0.67 and 0.28, and varimax 1.00, 0.94 and 0.61. The SAEs used several features for each strong lineage identity and none for most weak states. At depth 150 no method recovered the weak states, because information lost in the embedding cannot be recovered by any dictionary built on it.

**Figure 3.**
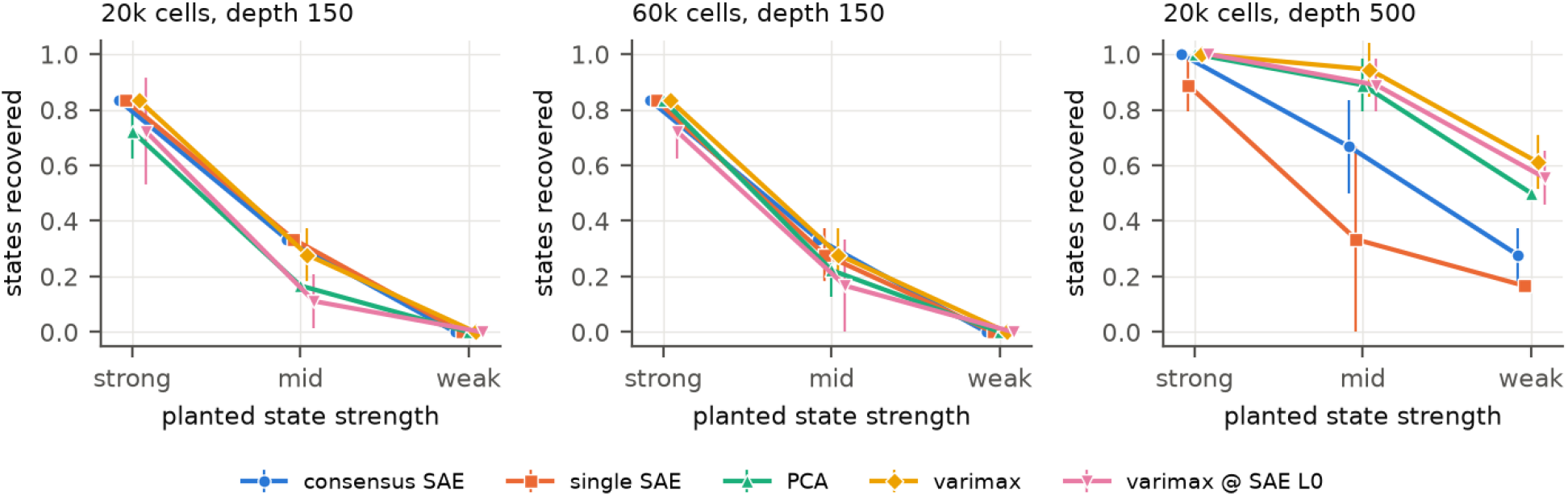
Recovery of planted cell states. Share of planted states recovered (r ≥ 0.7) by state strength, for three synthetic settings. At depth 150 no method recovers weak states. At depth 500 rotations recover most states, while single and consensus SAEs miss more of the mid-strength and weak states. Means and standard deviations over three seeds.

We then tested whether the discarded information could be recovered (Supplementary Tables S2 and S3). A single SAE trained for 15,000 steps, the total training of the five-fit consensus, did not reach the consensus reproducibility on any spatial embedding (0.42–0.67, against 0.85–0.96 for the consensus of the same SAE variant), and it lost most of its lineage-specific features as latents stopped activating (4–20, against 21–368 after 3,000 steps). A second SAE trained on what the consensus leaves unexplained, its residual, did no better. On planted data its consensus kept 4 of 50 features, which reproduced at 0.17 ± 0.29, and weak-state recovery rose only from 0.17 to 0.28, within the seed variation; on the CellCharter embedding of the second section, its features reproduced at 0.29 and covered no lineage. What the consensus discards is therefore mostly seed-specific detail rather than missed biology.

### The results do not depend on the SAE variant

Repeating all comparisons with a Matryoshka BatchTopK SAE^31^, which places coarse features first in the dictionary, gave the same results (Supplementary Table S1). On the spatial embeddings, single Matryoshka fits reproduced 24–45% of their features and their consensus 85–96%. The plain BatchTopK consensus was at least as reproducible (2 higher and 3 tied of 5 embeddings) and had slightly higher kNN accuracy (4 of 5), whereas the Matryoshka consensus produced fewer duplicate features per planted lineage (3.6 against 5.3).

## Discussion

Reproducibility and concept alignment measured different properties on every embedding we examined, and the reason lies in what each method optimizes. Varimax is a deterministic function of the covariance of the embedding, so it is reproducible whenever the covariance is estimated well, whatever its axes mean (PCA axes were less stable, because components of similar variance can exchange between halves of the data). Rotation axes are chosen for variance (PCA), simple loadings (varimax) or statistical independence (ICA), and none of these criteria ties an axis to a group of cells. Cell lineages in these embeddings are clusters rather than independent directions, so a dense rotation axis separates several lineages at once, which is what the chimeric gene programs of rotations show. A sparse code assigns each cell to a few active features and therefore favors features that follow cluster structure. This is why SAE features covered lineages with purer programs. Sparsifying varimax after fitting imposes the same constraint and recovered coverage on the PCA and Geneformer embeddings, but not purity: its axes were still fitted without it.

The same objective explains what SAEs miss. A reconstruction loss weights each direction by the variance it explains, and lineage identity dominates that variance. An SAE therefore spends several latents on each strong lineage and few on weak states; the duplicate identity features and the missed weak states are two consequences of one allocation. Orthogonal rotations cannot repeat a direction, so once the strong directions are taken, their remaining components reach weaker ones, which is why varimax recovered weak planted states that the SAEs missed. When weak states are the target, removing the dominant variance before fitting, for example by fitting within lineages, is a more direct solution than a larger dictionary; we did not test it here.

Seed dependence fits this picture. The SAE objective has many near-equivalent solutions, and strong concepts form the part of the dictionary that all of them share. A consensus over seeds keeps this shared part, as cNMF does for matrix factorization, so it is reproducible and as pure as a single fit. What it removes is mostly seed-specific: a longer single fit and a second-stage SAE on the residual both failed to recover new concepts, and the consensus held only modestly more reproducible features than one fit. Its value is mainly to mark which features can be trusted. This agrees with the finding in language models that features that vary between seeds still occupy reproducible low-dimensional subspaces^16^: the seeds differ in how they split a shared region, not in which regions they find.

How large this shared part is depends on the embedding. In the biological embeddings, most of the variance lies in a few directions (8.5–20 effective dimensions), and lineages alone explain 17–28% of it. Every seed finds these strong directions, and the consensus reached 85–98% reproducibility. The language-model embedding spreads its variance over about 70 effective dimensions, and token classes explain only 12% of it. Fits then have more ways to divide the space, fewer features are shared, and reproducibility reached 78%. A dictionary that is large relative to the number of cells has the same effect, which is why it had to be capped on Geneformer. A reproducibility value therefore reflects the embedding and the amount of data as much as the features, and a threshold set in one domain should not be carried to another.

These results bear on methods that make SAE features agree across seeds by construction, through aligned training, archetypal constraints or pooling^14–16^. Agreement is necessary for a feature to support a claim, but it can be obtained without concepts, for instance from shared initialization^17^ or, in the limit, from a fixed rotation. We suggest that such methods, and SAE analyses of biological embeddings generally, report reproducibility together with concept coverage and purity, against a sparsified rotation of the same embedding. Planted data are the cleanest benchmark, because they show both what no method recovers and how each method spends its features. The absence of a weak state from an SAE dictionary should not be read as its absence from the data.

Our study has limitations. Our ground truth is narrow: lineages, planted programs and rule-defined token classes, and a state that an embedding does not carry cannot be evaluated on it. We tested two SAE variants and models of up to 70 million parameters, on two spatial sections from one cancer type, which limits generalizability.

## Methods

### Data and embeddings

#### Spatial transcriptomic

We used two colorectal cancer sections profiled with the CosMx whole-transcriptome panel^20,21^. Section A (130,814 cells) is from a study of colorectal malignant transformation^22^; section B (493,834 cells) is the public CosMx Human Whole Transcriptome Colon Dataset (Bruker Spatial Biology). Both sections were analyzed on the same 2,000 genes (see Gene panel). From each section we sampled 50,000 cells uniformly at random (seed 0). Lineage labels came from supervised Insitutype cell typing^24^ against a colorectal cancer single-cell reference^25^, grouped into seven lineages (epithelial, fibroblast, myeloid, endothelial, mural, B/plasma, T/NK). The fine cell types were used for kNN accuracy. Embeddings were the first 50 principal components of log-normalized expression (log1p of counts per 10,000; Expr50), CellCharter^3^ latents (256 dimensions; both sections) and CadaST^23^ latents (256 dimensions; section B).

#### Gene panel

The 2,000 genes were selected by the preprocessing of our spatial pipeline in three steps. First, genes matching a regular expression for technical and ubiquitous gene families were removed: ribosomal proteins, mitochondrially encoded genes, translation elongation and initiation factors, poly(A)-binding proteins, ATP synthase and respiratory-chain subunits, mitochondrial translocases, and heat-shock and chaperonin genes. Second, marker genes of the cell types of the Pelka et al. reference^25^ (top differentially expressed genes per cell type) were included. Third, the panel was completed to 2,000 genes by a combined rank of expression dispersion (highly variable genes) and spatial autocorrelation (Moran’s I) computed over the sections. The list is provided with the code.

#### Single-cell RNA

We embedded 12,881 cells of the Pelka et al. colorectal cancer atlas^25^ with the Geneformer V2 model^2^ (1,152 dimensions). We tokenized each cell by Geneformer’s rank-value encoding, truncated to 2,046 genes, and averaged the penultimate-layer embeddings over gene tokens. Cells were sampled with at most 1,500 per lineage across nine reference lineages and scored over the same seven lineages as the spatial data. Gene programs were read on the genes shared with the CosMx panel.

#### Synthetic data

We simulated 1,000 genes and six lineages with proportions 0.35, 0.25, 0.15, 0.10, 0.10 and 0.05. Each lineage had a 50-gene identity program. It also had three state programs of 30 genes each, active in 35% of its cells with gamma-distributed usage and mixture weights 0.12, 0.06 and 0.03 (strong, mid and weak). Four shared programs (weight 0.06) were active in 15% of all cells. Expression rates combined a patient-shifted baseline (weight 0.45; four patients), the identity program (0.35) and the active programs. Counts were Poisson with log-normal library sizes at median depth 150 or 500. We used three settings (20,000 cells at depth 150 or 500; 60,000 cells at depth 150) and embedded each as Expr50.

#### Language model

We took residual-stream activations after layer 3 of Pythia-70m^26^ (512 dimensions) on the WikiText-103 corpus^27^, in 128-token windows. We discarded the first eight positions of each window to avoid attention-sink tokens. The main analysis used one million tokens from the training split: 900,000 for fitting and a held-out set of 100,000 for all metrics. A preliminary analysis used 50,000 validation tokens.

Tokens were assigned to eight classes by rule: numbers, punctuation, capitalized words, a closed list of function words, words ending in -ing, -ed or -ly, word-piece continuations, and “other” (excluded from scoring).

### Preprocessing

Each embedding was centered, each row was scaled to unit norm, and the whole matrix was divided by one scalar so that its mean squared entry was 1.

### Sparse autoencoders

We used BatchTopK SAEs^28^ with M latents. Training used tied initialization from k-means++ centers, unit-norm decoder rows, and a mean of k = M/16 active latents per row (k = 8 with M = 200 on synthetic data), selected as the k·B largest pre-activations in each batch of B rows. Inference used a global threshold calibrated so that the mean number of active latents on the training rows equals k. Dead latents received an auxiliary reconstruction loss (weight 1/32, 16 latents)^7^. Models were trained for 3,000 Adam steps^32^ with batch size 4,096 and learning rate 3 × 10^−3^, with warm-up and cosine decay, and small input noise early in training. The dictionary had M = 4d latents for embedding dimension d, capped at about one latent per 50 rows: 4d for the spatial (200–1,024 latents for 50,000 cells), synthetic (200 latents) and language-model embeddings (2,048 latents for 900,000 tokens), and 256 for Geneformer (12,881 cells). The Matryoshka variant^31^ adds reconstruction losses for the nested prefixes M/4 and M/2.

On Apple-silicon GPUs, a single top-k over about 1.8 × 10^9^ activations returned 0 without raising an error. For more than 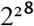 activations we therefore calibrate the threshold on the CPU from a 200,000-row sample. All smaller runs used the exact computation, and their sparsity matched the target.

### Consensus SAE

We fitted five SAEs with different seeds on the same rows. For every pair of fits, features were matched by the Hungarian algorithm^33^ on the Pearson correlation of activations, computed on up to 30,000 rows; matched pairs with r ≥ 0.7 were candidate links. Links were processed in order of decreasing correlation, and two groups of features were merged only if they contained no feature from the same fit and their mean cross-fit correlation was at least 0.7 (constrained average linkage). Each group thus holds at most one feature per fit, which prevents chains of pairwise matches from merging distinct concepts. Groups with features from at least three of the five fits were kept, and the consensus activation was the mean of the members’ activations. The rule treats all fits alike. For the anchored consensus (residual stage, Matryoshka consensus and Supplementary Table S5), features of the first fit were kept if matched at r ≥ 0.7 in at least two of the four other fits.

### Rotation baselines

PCA, varimax^18^ and FastICA^29^ were computed on the same preprocessed embeddings with d components. Each component was split into its positive and negative parts, giving 2d non-negative features. For sparse varimax, we kept each row’s k largest entries, with k equal to the mean number of active features of the consensus SAE in the same run and seed. On the language model, rotations were fitted on a 200,000-row sample of the fitting tokens.

### Two-stage residual model

Given consensus codes Z_1_ (centered) and the centered embedding X, we fitted a decoder D_1_ = (Z_1_^T^Z_1_ + λI)^−1^Z_1_^T^X, with λ equal to 10^−6^ of the mean diagonal of Z_1_^T^Z_1_. The residual R = X − Z_1_D_1_ is then uncorrelated with every column of Z_1_; we verified this numerically to within 10^−7^. The rescaled residual was fitted with a second SAE of 0.10M or 0.25M latents (k = M_2_/16), as a single fit or a consensus. The final code was the concatenation of the two stages.

### Metrics

#### Reproducibility (R1)

Rows were split into two random halves. The complete method (single fit, consensus, two-stage model or rotation) was fitted on each half, and both models encoded the same evaluation rows: all rows for the biological data, the held-out tokens for the language model. Features were matched by the Hungarian algorithm on activation correlation. R1 is the number of matched pairs with r ≥ 0.7 divided by the number of features of the smaller model.

#### Concept specificity and coverage

For each active feature, its concept selectivity is the largest share of its mean absolute activation in one concept, after averaging within concepts so that concept size does not matter. A feature is concept-specific if this share is at least 0.7. Coverage is the number of distinct concepts (lineages or token classes) with at least one concept-specific feature.

#### Gene programs, chimeras and Hallmark enrichment

For each feature, the gene program is the 50 genes with the largest regression slope of z-scored log-normalized expression on the z-scored activation. A feature is chimeric if its program contains at least three markers of lineages other than its own. Markers were derived from the single-cell reference^25^: up to 16 genes per lineage with mean expression at least four times higher in the lineage than in any other, detected in at least 20% of the lineage’s cells and in at most 30% of the cells of any other lineage. A feature counts as Hallmark-enriched if its program is enriched for an MSigDB Hallmark gene set^30^ (hypergeometric test against the panel; false discovery rate < 0.05).

#### Identity

For the biological data, kNN accuracy is the macro-averaged recall of fine cell types (with at least five cells) under leave-one-out 15-nearest-neighbor voting on the cosine distance of the full code. For the language model, it is the balanced accuracy of 15-nearest-neighbor classification of token classes, with five-fold cross-validation on 20,000 held-out tokens.

#### Planted programs

A planted program counts as matched if some feature correlates with its usage at r ≥ 0.7. Identity programs were evaluated over all cells, states within their lineage, and shared programs by the mean of their within-lineage correlations. Coverage on planted data is the number of matched programs (of 28). A feature counts as matching a program if its activation correlates at r ≥ 0.7 with at least one program over all cells. Identity duplicates are the features per lineage that correlate at r ≥ 0.7 with the lineage indicator.

#### Embedding geometry

The effective dimension is the participation ratio (Σλ_i_)^2^/Σλ_i_^2^ of the covariance eigenvalues λ_i_. The variance explained by concepts is the share of the total variance explained by the concept means.

### Statistics

Each configuration was run with three seeds, which set model initialization, the random half split and, for synthetic data, the simulation. We report means and standard deviations. In comparisons, a method wins on an embedding if its mean exceeds the other’s by more than the sum of their standard deviations; otherwise the comparison is a tie. Counts of wins, ties and losses are given per source.

### Software and compute

All models were implemented in PyTorch^34^ and trained on an Apple M-series GPU. Rotations, k-means++ initialization and kNN used scikit-learn^35^. Language-model activations were extracted with Hugging Face Transformers^36^. Evaluating one consensus SAE (three consensus fits: all rows and two halves, 15 SAEs in total) took 9–13 minutes on a spatial embedding, 12 minutes on Geneformer and about 39 minutes on the language model.

## Data availability

All data analyzed are public. Section A is section 232 of Crowell et al.^22^, available at Zenodo (10.5281/zenodo.15574384, file `232.h5ad`; CC-BY-4.0). Section B is the CosMx Human Whole Transcriptome Colon Dataset from Bruker Spatial Biology (https://brukerspatialbiology.com/products/cosmx-spatial-molecular-imager/ffpe-dataset/cosmx-human-whole-transcriptome-colon-dataset/)^21^. The colorectal cancer atlas^25^ is available from GEO (GSE178341). WikiText-103 and Pythia-70m are available from Hugging Face. Our derived data are deposited at Zenodo (https://doi.org/10.5281/zenodo.23672147): the cell identifiers, cell types and embeddings of the cells analyzed, the language-model activations, and the result files underlying every table and figure. Count data are available from the sources above.

## Code availability

Code for the SAE, consensus, rotation baselines, metrics, synthetic generator and figures, together with the list of the 2,000 genes, is available at https://github.com/AH-Shah/sae-groundtruth and archived at Zenodo (https://doi.org/10.5281/zenodo.23671822).

## Acknowledgements

We thank Amir Ata Arab Halvaei for methodological insights.

## Author contributions

All authors contributed equally to this work.

## Supplementary information

**Table S1.** Matryoshka SAE compared with the plain BatchTopK SAE. The Matryoshka consensus used the anchored rule.

|  | Matryoshka | Plain BatchTopK |
| --- | --- | --- |
| Consensus R1, spatial embeddings | 0.85–0.96 | 0.92–0.98 |
| Consensus R1, Geneformer (256 latents) | 0.80 | 0.85 |
| Single-fit R1, spatial embeddings | 0.24–0.45 | 0.31–0.44 |
| Plain vs Matryoshka consensus, R1 (W/T/L) | – | 2/3/0 |
| Plain vs Matryoshka consensus, kNN (W/T/L) | – | 4/1/0 |
| Identity duplicates per planted lineage (consensus) | 3.6 | 5.3 |

**Table S2.** Compute-matched single fit (Matryoshka SAE). The 15,000-step fit used one seed; 3,000-step fits and the consensus are means over three seeds.

| Embedding | R1, 15,000 steps | R1, 3,000 steps | R1, consensus | Lineage-specific features, 15,000 / 3,000 steps |
| --- | --- | --- | --- | --- |
| A, Expr50 | 0.42 | 0.45 | 0.96 | 4 / 25 |
| A, CellCharter | 0.67 | 0.36 | 0.93 | 7 / 328 |
| B, Expr50 | 0.42 | 0.40 | 0.85 | 5 / 21 |
| B, CellCharter | 0.50 | 0.32 | 0.91 | 8 / 368 |
| B, CadaST | 0.54 | 0.24 | 0.89 | 20 / 353 |

**Table S3.** Two-stage residual model. Stage 1 is the anchored consensus.

| Setting | Strong / mid / weak states recovered | R1 (combined) | R1 (second stage alone) | Second-stage features kept |
| --- | --- | --- | --- | --- |
| Planted, depth 500: anchored consensus only | 0.94 / 0.50 / 0.17 | 0.85 | — | — |
| + second-stage consensus, 10% of M | 0.94 / 0.50 / 0.28 | 0.82 | — | — |
| + second-stage consensus, 25% of M | 0.94 / 0.50 / 0.28 | 0.85 | 0.17 ± 0.29 | 3.7 ± 2.5 of 50 |
| + second-stage single fit, 25% of M | 0.94 / 0.50 / 0.28 | 0.40 | — | 50 |
| B, CellCharter (one seed), 10–25% of M | — | 0.89–0.92 | 0.29 | no lineage covered |

**Table S4.** Embedding geometry. Pythia-70m values were computed on 50,000 validation tokens.

| Embedding | Effective dimension | Variance in top 10 PCs | Variance explained by concepts |
| --- | --- | --- | --- |
| Pythia-70m, layer 3 | 70.5 | 0.29 | 0.12 |
| Geneformer V2 | 8.5 | 0.63 | 0.23 |
| Spatial embeddings | 10.0–19.7 | 0.46–0.66 | 0.17–0.28 |

**Table S5.** Anchored and symmetric consensus (three seeds; mean ± s.d.). Coverage and R1 as in Table 1; features = consensus features kept.

| Embedding | Coverage, anchored / symmetric | R1, anchored / symmetric | Features, anchored / symmetric |
| --- | --- | --- | --- |
| A, Expr50 | 2.0 / 2.0 | 0.98 ± 0.04 / 0.98 ± 0.03 | 39 / 40 |
| A, CellCharter | 3.0 / 3.3 | 0.92 ± 0.02 / 0.92 ± 0.02 | 399 / 444 |
| B, Expr50 | 3.0 / 3.0 | 0.92 ± 0.00 / 0.93 ± 0.04 | 39 / 40 |
| B, CellCharter | 4.3 / 5.0 | 0.92 ± 0.02 / 0.94 ± 0.01 | 351 / 377 |
| B, CadaST | 3.0 / 3.3 | 0.92 ± 0.02 / 0.93 ± 0.01 | 284 / 319 |
| Planted, 20k cells, depth 150 | 16.3 / 17.0 | 0.88 ± 0.05 / 0.92 ± 0.07 | 31 / 30 |
| Planted, 20k cells, depth 500 | 19.7 / 21.7 | 0.85 ± 0.02 / 0.95 ± 0.04 | 43 / 48 |
| Planted, 60k cells, depth 150 | 16.3 / 17.0 | $0.87 \pm 0.05$ / $0.89 \pm 0.07$ | 32 / 34 |
| Geneformer | 7 / 7 | $0.81 \pm 0.01$ / $0.85 \pm 0.02$ | 130 / 141 |
| Pythia-70m | 8 / 8 | $0.78 \pm 0.01$ / $0.78 \pm 0.00$ | 601 / 690 |

